# Projected impacts of climate change on plant-frugivore interactions across the Americas

**DOI:** 10.1101/2025.03.05.641042

**Authors:** Alexandre Rabeau, Alexander Pigot, Joseph Tobias, Matthias Schleuning

## Abstract

**Aim:** Frugivorous animals provide crucial seed-dispersal functions in terrestrial ecosystems. However, the potential effects of climate change on the interactions between plants and frugivores are largely unknown. In this study, we map plant-frugivore interactions across the Americas and then project potential changes in this interaction diversity under future scenarios of climate change.

**Location:** North, South and Central America.

**Time Period:** Present (1980–2010) and future (2070–2100).

**Major Taxa Studied:** Frugivorous birds and fruit-bearing plants.

**Methods:** Based on a trait-matching framework, we estimated interaction probabilities between frugivorous birds (n = 629 species) and fruit-bearing plants (n = 3283 species) to construct ecoregion-level plant-frugivore networks. To estimate future climate impacts, we compared the climatic niches of birds with ecoregion climates and weighted interaction probabilities based on the overlap between avian climatic niches and current and future ecoregion climates. Lastly, we used simulations to examine how avian dispersal and dietary flexibility may buffer the future diversity of plant-frugivore interactions.

**Results:** Under future climate scenarios, our models projected a decline in interaction diversity across most ecoregions, especially in tropical and subtropical ecoregions and biomes. In addition, simulations indicated that interaction rewiring through dispersal and dietary flexibility had the potential to mitigate future losses of interaction diversity, particularly in tropical forests.

**Main conclusions:** Our results suggest that plant-frugivore interactions are at risk of losing diversity under continued rapid climate change, which may undermine their stability and resilience. However, projected losses of interactions depend on the capacity of frugivorous bird species to modify their ranges or dietary preferences in future. As it is unknown to what extent these factors can buffer the stability of future interaction networks, the maintenance of high plant-frugivore interaction diversity in current networks is crucial to buffer ecosystems against future climatic changes, especially in the tropics.

## INTRODUCTION

The composition of ecological assemblages, and thus the potential for interactions among coexisting species, is changing rapidly in response to climate change (Scheffers et al., 2016). Some species are declining as conditions change, leading to local or global extinction (Román-Palacios & Wiens, 2020; Pecl et al., 2017). Other species survive by moving across landscapes as they track their climatic niche, leading to reshuffled species assemblages and novel interactions (Gallagher et al., 2013; Nolan et al., 2018). Given that a wide range of mutualistic and antagonistic interactions between species play a fundamental role in ecological processes, such as predation, pollination and seed dispersal, it is clear that climate change may have serious consequences for the resilience of ecosystem functions and services derived from these interactions (Díaz et al., 2018). Nonetheless, the impacts of future climate change on species interactions are largely unknown at macroecological scales.

The interactions between fruit-bearing plants and frugivorous animals is an essential process to many ecosystems (Lim et al., 2020; Howe & Smallwood, 1982), particularly in the tropics with 60-80% of all plant species dependent on frugivores (Jordano et al., 2011). Animal-mediated seed dispersal underpins forest regeneration, fostering plant functional connectivity and resilience to habitat degradation (McConkey et al., 2012; Mueller et al., 2014). Additionally, frugivory by mobile animals provides a key avenue for plant responses to climate change as the dispersal of seeds across the landscape helps plants to track their climatic niche (Sales et al., 2021), which can require movements of up to 10km/year (González-Varo et al., 2017). Despite the importance of frugivory and the sensitivity of plant-frugivore interactions to environmental change (Neuschulz et al., 2016), climate change impacts on plant-frugivore interactions are less known than that of other mutualistic interactions such as pollination (Teixido et al., 2022). It is therefore essential to investigate how the diversity of plant-frugivore interactions is likely to respond to projected climate change.

Studies that have investigated climate change impacts on plant-frugivore interactions have generally been restricted to small taxonomic (Mackay & Gross, 2019; Sales et al., 2021) and geographic scales (Bender et al., 2019; Dehling et al., 2016). Previous large-scale studies have mostly compared patterns in species richness between specific groups of prospective interaction partners (Kissling et al., 2007; Mota et al., 2022). Another widely used method is the construction of interaction networks from observed interaction data (Maglianesi et al., 2014; Quitián et al., 2019). Whilst this method is powerful to describe species interactions for specific ecological assemblages, site specific biases and constraints in data availability make it difficult to utilise such empirical data at large scales (Kissling & Schleuning, 2015). To overcome such limitations, previous studies have developed trait-based models to estimate interaction probabilities between plants and birds at macroecological scales (Fricke et al., 2022; Nowak et al., 2022). These models build on the matching between plant and bird morphology, especially between fruit size and avian gape width (Dehling et al., 2016; Wheelwright, 1985). We combine this method with climate niche estimates for frugivorous bird species to investigate potential impacts of climate change on plant-frugivore interactions at geographic and taxonomic scales previously not accessible.

The interactions between plants and birds are dynamic and are likely to change in response to environmental change (Tylianakis & Morris, 2017). First, many species are expected to shift their ranges in response to climate change to track their climatic niche (Schleuning et al., 2020). Second, species have shown a great capacity to adapt their diet and interact with a wider or different range of resources if resource availability changes (Bender et al., 2017). Both these mechanisms can lead to “interaction rewiring” – the formation of novel interactions between species – either because novel interactions form with immigrating species or because presently co-occurring species form new interactions (CaraDonna et al., 2017; Schleuning et al., 2020). Integrating such mechanisms of interaction rewiring into simulations is important to assess the potential vulnerability of plant-frugivore interactions to future climate change.

In this study, we use a trait-based approach (Figure 1) to assess projected impacts of climate change on plant-frugivore interactions at a macroecological scale across North and South America. Our aim is to first derive patterns in plant-frugivore interaction diversity – the number of avian interaction partners potentially available to a plant species – across the Americas. Given historical climate patterns driving evolutionary radiations in vascular plant and subsequent frugivore diversity, we hypothesize plant-frugivore interaction diversity to increase towards the tropics (Kissling et al., 2009). Second, we aim to determine the exposure of fruit-bearing plants to climate change by assessing the suitability of future climate to their avian interaction partners (Figure 1). At lower latitudes, we hypothesize plant-frugivore interaction diversity to decrease in response to climate change, as we expect future climate to be less suitable for frugivorous birds than current climate, especially as bird species inhabiting these regions may have narrower dietary breadths and climatic niches (Cadena et al., 2012; Dalsgaard et al., 2017; Mota et al., 2022). Conversely, at higher latitudes, we expect potential increases in interaction diversity as climatic conditions become more suitable for avian frugivore partners (P. M. Martins et al., 2024). Finally, we aim to understand the extent to which interaction rewiring may influence projections of plant-frugivore interactions under future climate (Figure 1). Clearly, any such future projection is complicated by the fact that losses in interaction diversity can potentially be compensated by the emergence of novel interactions due to avian dispersal and/or dietary flexibility (CaraDonna et al., 2017; Schleuning et al., 2020). The direction of such effects is difficult to predict because, on one hand, avian dietary flexibility and dispersal ability peak at high latitudes (Morelli et al., 2021; Sheard et al., 2020), potentially increasing the number of species able to buffer any local losses in interaction diversity, whereas, on the other hand, the number of potential replacement species and dietary options are much higher in the tropics (Kissling et al., 2009; Schleuning et al., 2012). We attempt to disentangle these two possibilities by accounting for dispersal ability and diet breadth in our models.

**Figure 1.**
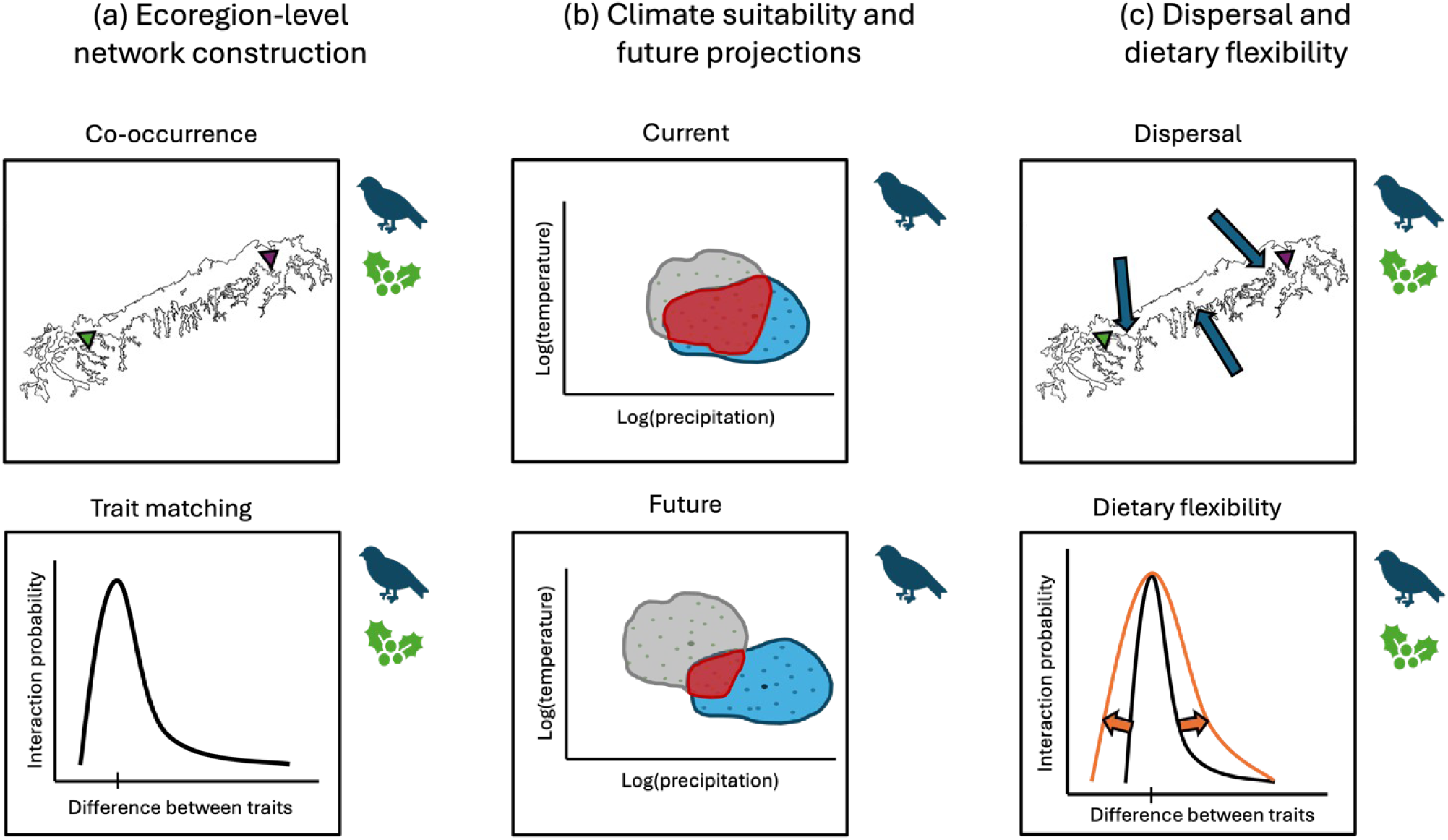
Projecting climate change impacts on plant-frugivore interactions. (a) We construct ecoregion-level networks by first establishing co-occurrence between plant and bird species (top). Co-occurrence is established if a given bird (blue triangle) or plant (green triangle) species both occur within the same ecoregion. Interaction probabilities between prospective interaction partners are then calculated from a trait-based interaction model (bottom). As shown, interaction probability increases as the difference between fruit and gape size approaches 0. (b) Climatic suitability of an ecoregion for avian frugivores was estimated by calculating the level of overlap (red) between the avian climatic niche hypervolume (blue) and the ecoregion climate hypervolume (grey) under current and future climatic conditions. If the Sorensen’s overlap was below 0.4, the bird species was dropped out of the community in this ecoregion. (c) We accounted for potential positive effects of avian dispersal (top) and dietary flexibility (bottom) in future projections of plant-frugivore interactions. To integrate dispersal, we allowed species from surrounding ecoregions to immigrate into ecoregion networks. To integrate dietary flexibility, we modified the trait matching parameter from a tighter relationship (black) to a looser relationship (orange) between fruit size and gape width.

## METHODS

To assess the effects of climate change on plant-frugivore networks, we projected current and potential future patterns in plant-frugivore interaction diversity by using a simulation model with three consecutive steps (Figure 1). First, we constructed plant-frugivore networks at an ecoregion-level using a trait-based interaction model informed by functional trait data of co-occurring plant and frugivorous bird species. Second, we determined the indirect exposure of plant species to climate change by comparing the climatic niche of their respective avian interaction partners with current and future climate. Third, we quantified how interaction rewiring via avian dispersal into ecoregions and increased dietary flexibility may modify projected future patterns in plant-frugivore interaction diversity.

Our main analysis included obligate avian frugivore species with at least 50% of fruit in their diet (Wilman et al., 2014), corresponding to the group of frugivores that are the main seed dispersers in seed-dispersal networks (Schleuning et al., 2014). In an additional analysis, we lowered this threshold to 25% (Wilman et al., 2014) to include opportunistic frugivores which only occasionally feed on fruits and can be important seed dispersers, especially in marginal habitats and/or at higher latitudes (Dalsgaard et al., 2017; Schleuning et al., 2011). Hence, we ran all analyses once for obligate frugivores with >50% of fruit diet (n = 629 species) and once for obligate and opportunistic frugivores with >25% of fruit diet (n = 903 species).

### Distribution data

To establish plant and bird species distributions, we used comprehensive range map datasets for the two groups. For fruit bearing plants, we extracted range map data from the BIEN database (Maitner et al., 2018). The BIEN database is composed of range map estimates for over 98,000 New World Plants, generated primarily through the implementation of Maximum Entropy distribution modelling informed by over 81 million georeferenced occurrence records (Maitner et al., 2018). In the case of birds, we extracted expert range maps from BirdLife (BirdLife International and Handbook of the Birds of the World, 2021). The Birdlife Species Range map database contains range map estimates for >11,000 bird species globally, derived by experts from over 5.6 million georeferenced occurrence records (BirdLife International and Handbook of the Birds of the World, 2021). We subset this dataset to only include resident, wintering and breeding ranges of birds. Range map datasets are particularly effective at describing broad regions where a species is likely to occur, particularly for widespread species (Merow et al., 2017). However, a key limitation is that these datasets tend to overestimate species occupancy (Merow et al., 2017).

### Trait data

We compiled functional trait data for plants and birds, focusing on traits providing information about trophic ecology, particularly fruit and beak dimensions. For fruit-bearing plants, we compiled a dataset from several sources providing data on fruit dimensions for 3,283 plant species that occur across the Americas (Dehling et al., 2016; Kissling et al., 2019; Sinnott-Armstrong et al., 2018). We included plant species primarily dependent on birds for seed dispersal by filtering for plants with contrastive fruit colour syndromes (Sinnott-Armstrong et al., 2021). Fruit length (mm) was the plant functional trait with the highest coverage and is closely correlated with other fruit dimensions (McFadden et al., 2022). To inform our trait-based interaction model, we used fruit length rather than other plant functional traits such as seed size as fruit length has been shown to correlate strongly with avian gape width and would therefore more accurately determine interaction probability (Bender et al., 2018; Dehling et al., 2014). Given that most frugivorous birds consume the whole fruit, a key trait constraining or predicting plant-bird interactions is gape size (Wheelwright, 1985). We extracted gape-size (mm) for 629 frugivorous bird species from McFadden et al. (2022) and Tobias et al. (2022), which was measured as the horizontal distance between the points where the lower and upper mandibles connect.

### Determining bird and plant co-occurrence at ecoregion level

We established co-occurrences between fruit-bearing plant and frugivorous bird species to understand which pairs of species could potentially come into contact based on their current distributions. To do so, we first divided our study system into terrestrial ecoregions (n = 277) (Olson et al., 2001). Ecoregions have been shown to be a strong delineation for interacting plant-frugivore assemblages (L. P. Martins et al., 2022). We constructed co-occurrence matrices by intersecting each plant and bird species’ range map with that of a given ecoregion. These assemblages thus represent the pool of plant and bird species that could potentially interact at the ecoregion level. For each ecoregion-level assemblage, we mapped the median and variance of fruit length, gape width and range size across all plant and bird species, respectively (Figure 2a-d) (Figure S1).

**Figure 2:**
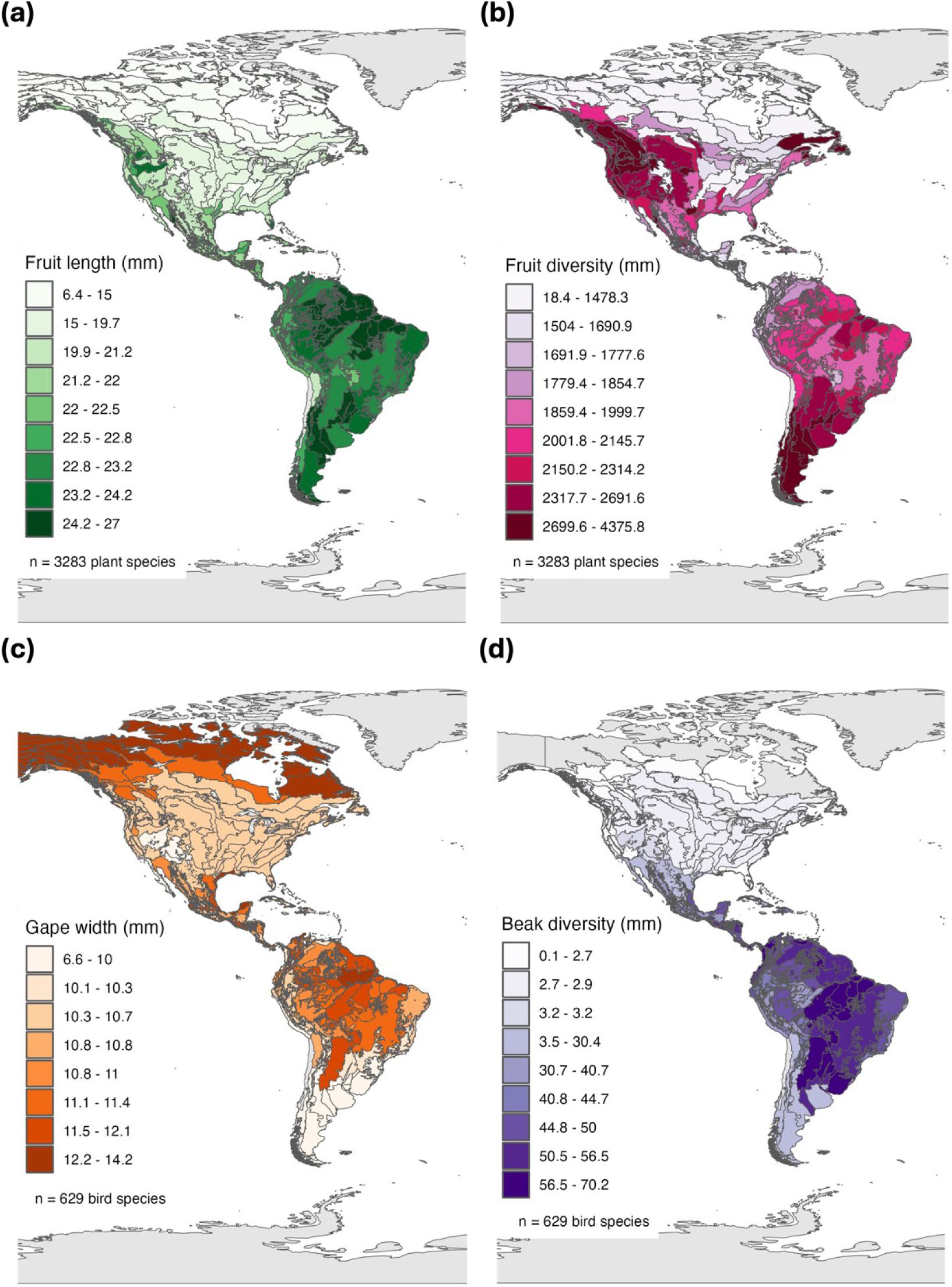
Latitudinal patterns in fruit length and avian gape width across the Americas. (a) Map of median fruit length across the Americas. Regions shaded in darker green are occupied by plant communities with high median fruit length. (b) Map of fruit length variance across the Americas. Regions shaded in darker pink are occupied by plant communities with a high variance in fruit length. (c) Map of median gape width of obligate frugivores across the Americas. Regions shaded in darker orange are occupied by bird communities with high median gape width. (d) Map of gape width variance of obligate frugivores across the Americas. Regions shaded in darker purple are occupied by bird communities with a high variance in gape width.

### Climate data

To describe the range of climatic conditions tolerable by each frugivorous bird species, we generated climatic niche hypervolume based on average monthly precipitation (10g/m^2^) and temperature (°C) for processed eBird occurrence records at a 1 km resolution and at the month and year of the occurrence (Karger et al., 2017; Sullivan et al., 2009). Temperature and precipitation (log transformed) were selected as climate variables to inform niche estimates as they have previously been shown to strongly shape avian distributions (Pigot et al., 2010). Niche hypervolumes were generated using the box kernel density method, with the Silverman bandwidth estimator used to provide an appropriate bandwidth (Blonder et al., 2014). We used a consistent method to summarize present and future climatic conditions for a given ecoregion. For current climate, average monthly temperature and precipitation layers (1 km resolution, 1980 – 2010) were extracted using the ecoregion polygon boundaries (Karger et al., 2017) and used to generate hypervolumes summarizing the multivariate range of climates within each given ecoregion (Blonder et al., 2014). This method thus used an identical spatial and temporal resolution of climate data used in estimating avian climate niches. For projected future climate (2070–2100), the same method was implemented across four climate models (GFDL-ESM4, IPSL-CM6A, MPI-ESM1, MRI-ESM2) under an RCP 7.0 emissions scenario which corresponds closest to ongoing rapid temperature increase (Pachauri R.K. & Meyer L.A., 2014).

### Trait-based interaction model

To determine plant-frugivore interaction diversity across the Americas, we constructed probabilistic plant-frugivore networks at the ecoregion level using a trait-based model. With this model, it was possible to derive an interaction probability between a bird and plant species depending on the level of trait correspondence – determined as fruit size (mm) subtracted from gape width (mm) (Figure 1). The relationship between trait correspondence and interaction probability followed a theoretically determined right-skewed shape as birds with larger beaks are able to interact with smaller fruit, while small birds are unlikely to access larger fruits (Nowak et al., 2022; Wheelwright, 1985).

In the model, the degree of network specialization was adjusted through a trait matching parameter (s) (Donoso et al., 2017). Modifying this parameter either widens or tightens the range of resources a bird can interact with. To represent a realistic degree of specificity comparable to empirical plant-frugivore networks, we set this parameter to s = 1.5 to construct the networks under current conditions (Nowak et al., 2022). This s-value corresponded to a mean complementary specialization of H_2_’ = 0.43 ± 0.11 in the ecoregion-level networks (Schleuning et al., 2012). A high H_2_’ number determines a high level of specialization within an interaction network – i.e. most interactions occur between a fewer number of interaction partners - whereas a low H_2_’ number is indicative of more generalism within an interaction network (Blüthgen et al., 2006).

### Current and future climatic suitability

To determine the impact of projected climate change on plant-frugivore interactions, we assessed the climatic suitability of current and projected future ecoregion climate for avian frugivores. For a given network, the climatic niche of each avian frugivore was compared to the ecoregion climate. Subsequently, every interaction probability between a given plant and bird species in an ecoregion was weighted by the climatic suitability of the respective ecoregion. The climatic weighting was calculated as the Sorensen’s overlap (range: 0-1) between the hypervolumes of the avian climatic niche and the current and future ecoregion climate, respectively. A high overlap between the avian niche hypervolume and the ecoregion climate hypervolume indicates the ecoregion is climatically suitable resulting in a high interaction probability between a bird and its plant partners. In contrast, a low overlap between the avian niche hypervolume and the ecoregion climate hypervolume indicates the ecoregion is climatically less suitable and will result in a lower interaction probability, for instance due to low abundance, local extinction or emigration of the bird species from that ecoregion (Thomas et al., 2006). We excluded bird species from the ecoregion community if only a fraction of the respective avian niche hypervolume overlapped with the ecoregion niche hypervolume. We assume here that a bird species is unlikely to occur in ecologically relevant numbers in an ecoregion that is mostly climatically unsuitable as the species would either go locally extinct or emigrate. For the main analyses, we set this overlap threshold to 0.4 – below which the respective bird species is excluded from the ecoregion community - however we also tested different thresholds (Figure S2). By applying the climatic suitability weighting method according to both current and future climate conditions, we used this approach to quantify projected impacts of changing climatic conditions on plant-frugivore interactions.

### Projections of interaction diversity

To determine projected changes in plant-frugivore interaction diversity, we first calculated the potential number of interaction partners for each plant species in an ecoregion based on the Shannon diversity of weighted pair-wise interaction probabilities (Dormann et al., 2009). We chose this metric because it measures the potential number of bird partners available to each plant species (Dormann et al., 2009). The same metric was calculated before climate weighting (Figure 3) and accounting for both current and projected future climatic conditions. We averaged interaction diversity across all plant species occurring in an ecoregion. To compare current and projected future interaction diversity, we calculated the proportional decrease or increase in the number of avian partners per plant species averaged both across all plant species in a given ecoregion and all the four climate models. Additionally, given the importance of frugivory in forests (Côrtes & Uriarte, 2013), we compared the proportional changes in interaction diversity across forest biomes by averaging ecoregion-level values for a given forest biome (Olson et al., 2001).

**Figure 3.**
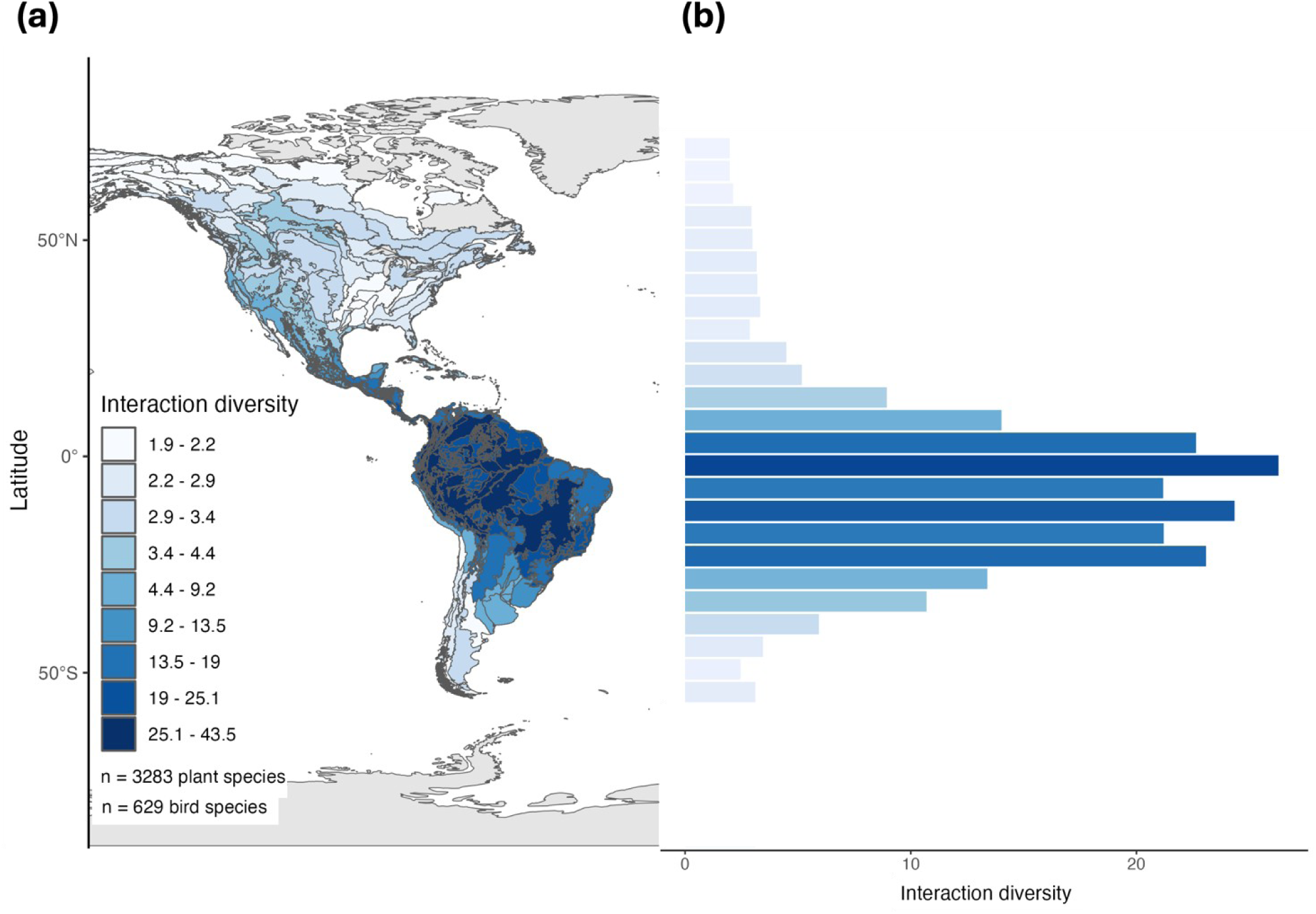
Latitudinal pattern in potential plant-frugivore interaction diversity across the Americas. Map of mean interaction diversity within an ecoregion (a) and across 5° latitudinal bands (b). Interaction diversity is defined as the predicted number of obligate avian frugivores per plant species. Darker blue ecoregions are occupied by plant species that, on average, are able to interact with many more frugivorous birds compared to plant species that occupy lighter blue ecoregions. Plant interaction diversity shown here is not weighted by the climatic suitability of the respective ecoregions. Projections including both obligate and opportunistic frugivores are shown in Figures S3.

### Interaction rewiring

We investigated the effects of interaction rewiring on future plant-bird interaction diversity by allowing for avian dispersal into an ecoregion and varying frugivore dietary flexibility. First, we simulated the dispersal of bird species into ecoregions, potentially offering compensation through interaction rewiring with immigrating species (CaraDonna et al., 2017; Schleuning et al., 2020). We calculated maximum dispersal distances for each bird species using a method adapted from Stewart et al. (2020). This involved fitting a linear mixed effects model between avian functional traits and empirical maximum natal dispersal distances for a subset of the bird species (n = 285 species) and then using the fitted model to predict maximum dispersal distances for the species used in this study (Stewart et al., 2020). All possible combinations of linear and quadratic effects for hand-wing index, wing length, generation length, body mass and range size were fitted against maximum dispersal distance as these features have been shown to affect dispersal distance (Stewart et al., 2020). The model with the lowest AIC score was a function of the linear effects of wing length, hand-wing index, geographical range size and a quadratic effect of hand-wing index (marginal R^2^ = 0.43, random terms = bird order/family). A dispersal buffer zone was then derived for each species by multiplying the predicted maximum dispersal distance by the number of generations that could occur between the midpoint of our current (1995) and future climate periods (2085) (Bird et al., 2020). The established ecoregion assemblages were then supplemented with those species for which the minimum distance between the source and destination ecoregion borders was less than the dispersal buffer. While this would allow species to migrate into climatically unsuitable ecoregions, we accounted for this by down-weighting interaction probabilities and excluding bird species from ecoregions with unsuitable climates (see above). Given established ecological barriers between biogeographic realms, species were only allowed to enter other ecoregions if both source and destination ecoregions belonged to the same realm (Stewart et al., 2020). For instance, a bird species originating from a Nearctic ecoregion could enter another Nearctic ecoregion but wouldn’t be able to enter any Neotropical ecoregions. Nevertheless, this is an optimistic scenario as species would be able to immigrate into new ecoregions but would not emigrate unless the climatic suitability drops below a threshold (see above). To quantify the effect of avian dispersal on the projected future plant-frugivore interactions, we reran the analysis based on the newly assembled ecoregion assemblages which included the immigrated bird species.

Second, to reflect the capacity of species to adapt their diet as resource availability changes, we allowed avian interaction partners to adapt to a more generalist fruit choice by varying the specialization parameter (s) in the trait matching model (Bender et al., 2017; Donoso et al., 2020). We therefore modified the s parameter of our trait-based interaction models from moderate (s = 1.5) to generalized (s = 0.5) trait matching (Nowak et al., 2022), resulting in a more generalized scenario corresponding to a mean complementary specialization of H_2_’ = 0.25 ± 0.05 at the ecoregion level. A lower specialization parameter allows frugivorous birds to form new interactions with other fruit-bearing plants, due to a relaxed trait matching between fruit size and gape width. To quantify the effect of increased dietary flexibility on projected future plant-frugivore interactions, we reran the analysis using this relaxed trait matching with and without avian dispersal into an ecoregion.

## RESULTS

### Current patterns in trait diversity and plant-frugivore interaction diversity

We found trait-specific latitudinal patterns in fruit size and gape width across the Americas. Median fruit length was highest across the Neotropical realm and in a few ecoregions on the west coast of North America (Figure 2a). The most diverse ecoregions, with regards to the variance in fruit length, were located around Patagonia and western North America (Figure 2c). Median gape width of obligate frugivores was highest at lower latitudes across Central America and Amazonia. Variance in gape width was highest in the Neotropics, with particularly high diversity of gape width in Eastern and Northern Amazonia (Figure 2d).

As expected, we found a strong latitudinal gradient in plant-frugivore interaction diversity and an increase in the mean number of avian interaction partners available to plants towards the tropics (Figure 3). For instance, the mean number of obligate frugivores potentially available to plant species was 38 species in the Southwest Amazon moist forests. Conversely, plant species occupying ecoregions at higher latitudes had much fewer potential interaction partners. For instance, in the Low Arctic Tundra, the average number of obligate frugivores per plant species was only two species. In the analysis including obligate and opportunistic frugivores, the latitudinal patterns remained qualitatively similar, but plant species at high latitudes had a higher number of potential avian interaction partners (Figure S3).

### Projected future changes in plant-frugivore interaction diversity

Future climate projections revealed widespread declines in the number of potential avian interaction partners across most of the Americas, both in the analysis of obligate frugivores (Figure 4a) and that including obligate and opportunistic frugivores (Figure S4a). For obligate frugivores, the model projected a decline in future interaction diversity for 71% of the 277 ecoregions. The highest declines in plant-frugivore interaction diversity were concentrated across Central Amazonia (projected decrease in interaction diversity: 100%), as well as the eastern United States (projected decrease: 100%). For a minority of ecoregions (n = 70), mostly those occupying high latitudes, the simulations of obligate frugivores projected a slight increase in the potential number of avian interaction partners as in these ecoregions climatic conditions largely became more suitable for avian species. Across the forest biomes where avian seed dispersal is particularly relevant, we found a consistent decline in projected plant-frugivore interaction diversity under projected future climate, particularly in tropical and subtropical ecoregions and biomes (Figure 4b, Figure S4b). For obligate frugivores, the highest declines in interaction diversity occurred in Mangroves (mean projected decrease: 37%). The analysis including both obligate and opportunistic frugivores showed qualitatively similar trends and a significant decline in projected interaction diversity across the tropical forest biome (Figure S4b).

**Figure 4.**
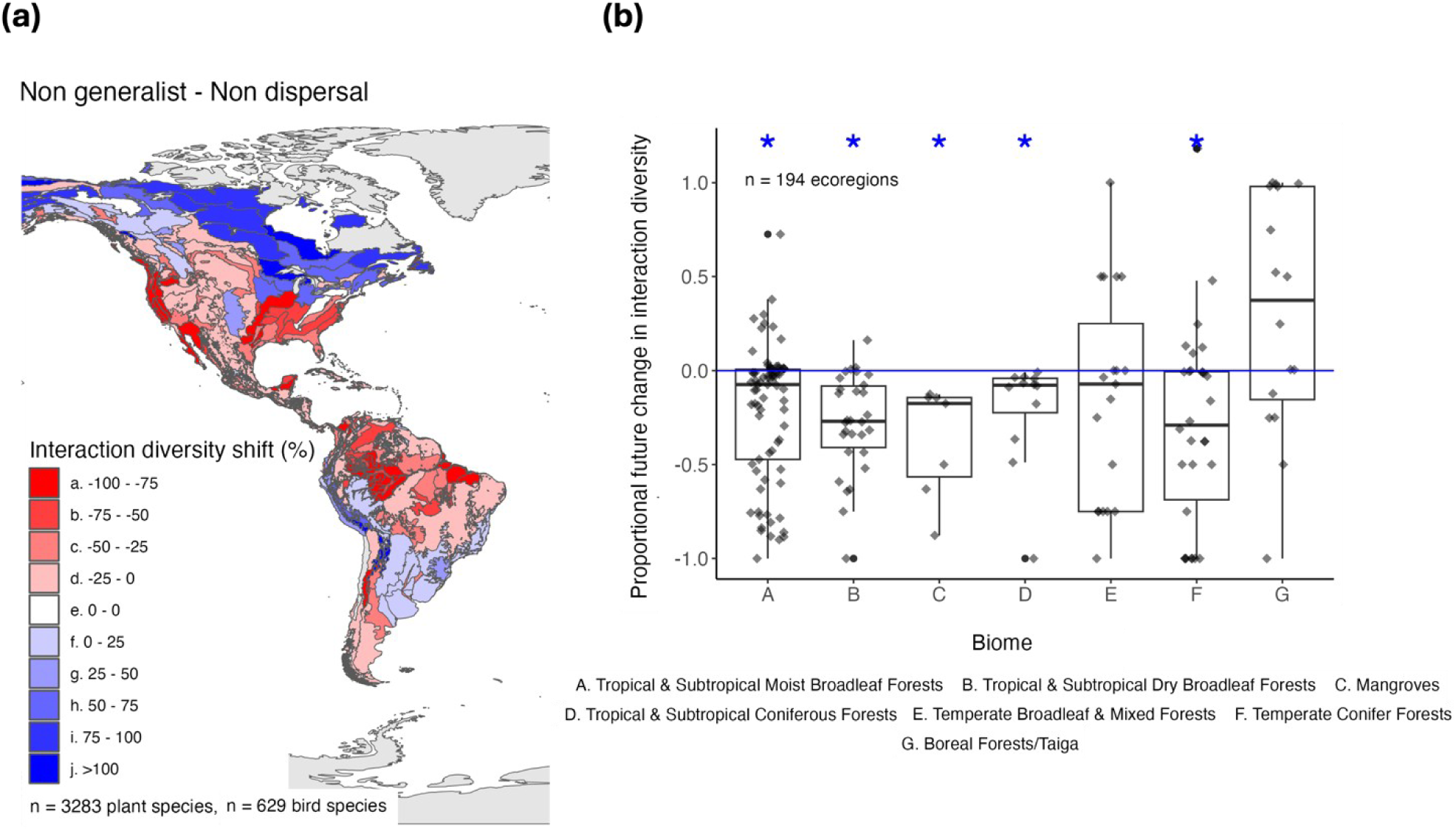
Projected impacts of climate change on plant-frugivore networks. (a) Map of future climate impacts on plant-frugivore networks. (b) Boxplot of future climate impacts on plant-frugivore networks. Proportional change in interaction diversity was calculated by dividing future number of interaction partners by current number of interaction partners and substituting by 1. Asterisks above each boxplot indicate significant differences from 0 based on one-sample t-tests. Shown are simulation results from the projections including only obligate frugivores. Projections including both obligate and opportunistic frugivores are shown in Figure S4.

### Effects of interaction rewiring on projected future plant-frugivore interaction diversity

Simulations of obligate frugivores that allowed for increased frugivore dietary flexibility and dispersal into ecoregions showed widespread positive effects on plant-frugivore interaction diversity across South and Central America, whereas interaction rewiring had weaker effects on the projected changes in plant-frugivore interaction diversity across North America. In the simulations allowing for increased dietary flexibility, 58% of the 277 ecoregions exhibited an increase in the interaction diversity, particularly around Western Amazonia and Central America (Figure 5a). In simulations allowing for dispersal, 63% of the 277 ecoregions exhibited an increase in projected plant-frugivore interaction diversity, with the strongest positive responses across Central and South American ecoregions, particularly in Patagonian and eastern Amazonian ecoregions (Figure 5b). Once we combined the effects of increased dietary flexibility and dispersal, a positive effect was exhibited in 70% of the 277 ecoregions, indicating a widespread increase in projected interaction diversity across Neotropical ecoregions (Figure 5c). Increased dietary flexibility and dispersal resulted in a consistent increase in projected interaction diversity in all tropical forest biomes (Figure 5). Simulations of obligate and opportunistic frugivores that allowed for increased dietary flexibility and/or dispersal showed widespread positive effects on interaction diversity across both North and South America (Figure S5). In the scenario allowing for both increased dietary flexibility and dispersal, 86% of the ecoregions showed an increase in the projected number of obligate and opportunistic frugivores under future conditions. In contrast to the simulations restricted to obligate frugivores, increased dietary flexibility and avian dispersal resulted in a strong positive increase in interaction diversity in both temperate and tropical forest biomes (Figure S5).

**Figure 5:**
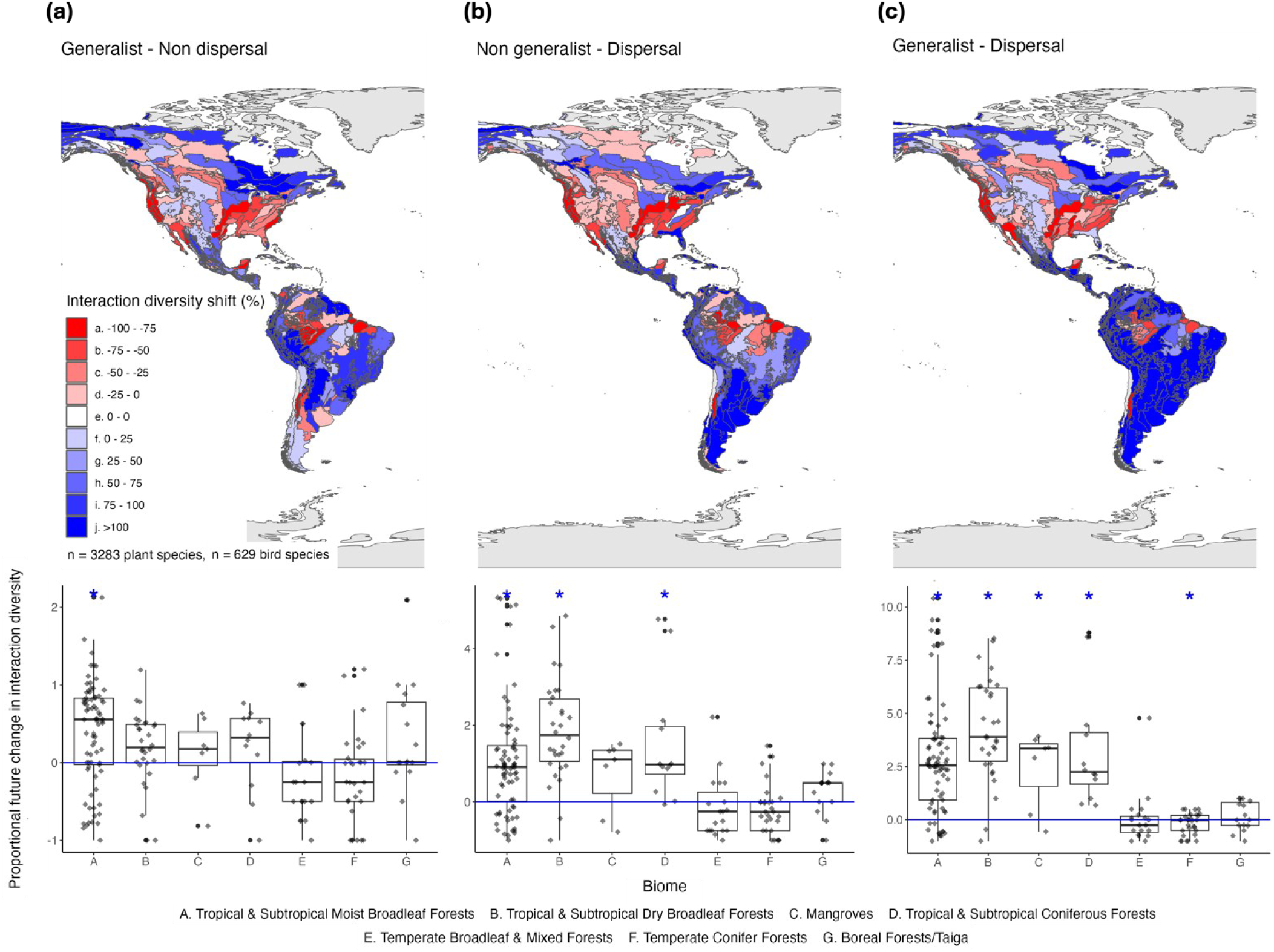
Effects of interaction rewiring on projected future plant-frugivore networks. (a) Map and boxplot of future climate impacts on plant-frugivore networks allowing for dietary flexibility (i.e., trait matching parameter *s* set to 0.5). (b) Map and boxplot of future climate impacts on plant-frugivore networks allowing for avian dispersal (i.e., immigration of species from surrounding ecoregions). (c) Map and boxplot of future climate impacts on plant-frugivore networks allowing for both avian dispersal and dietary flexibility. Proportional chance in interaction diversity was calculated by dividing future number of interaction partners by current number of interaction partners and substituting by 1. Asterisks above each boxplot indicate significant differences from 0 based on one-sample t-tests. Shown are simulation results from the projections including only obligate frugivores. Projections including both obligate and opportunistic frugivores are shown in Figure S5. Note the different scaling on the y-axes for the different scenarios and the additive effects of dietary flexibility and dispersal on changes in interaction diversity.

## DISCUSSION

We used a trait-based interaction model based on functional trait and range data of co-occurring fruit-bearing plants and frugivorous birds to project plant-frugivore interaction diversity across the Americas. Under projected future climate, our simulations suggest widespread and consistent declines in plant-frugivore interaction diversity across most ecoregions, especially in tropical forest biomes. However, both avian dispersal and increased dietary flexibility had the potential to increase plant-frugivore interaction diversity, especially in the Neotropics.

### Current patterns in trait diversity and plant-frugivore interaction diversity

Our models predicted geographic patterns in trait and interaction diversity that are consistent with empirical studies (Chen et al., 2017; Moles et al., 2007). Increased diversity of both fruit-bearing plants and frugivorous birds at lower latitudes may be driven by the enhanced productivity in these ecosystems (Kreft & Jetz, 2007; Ting et al., 2008). Furthermore, major radiations in fruit-bearing plant diversity in South and Central America may explain subsequent increases in frugivorous bird richness and trait diversity (Figure 3), particularly in the Passeriformes (Kissling et al., 2009). Our predictions are also in line with previous macroecological studies that have identified the Neotropics as the most significant hotspot with regards to plant-frugivore interaction diversity (Kissling et al., 2009). This latitudinal pattern in frugivore diversity was most pronounced for obligate frugivores whereas the diversity of obligate and opportunistic frugivores declined less steeply from Neotropical to Nearctic ecoregions (Kissling et al., 2009). Our trait based interaction model therefore accurately captures the current biogeographic patterns in plant-frugivore interactions.

### Projected future changes in plant-frugivore interaction diversity

Our simulations projected widespread reductions in plant-frugivore interaction diversity under future climate, especially in tropical and subtropical ecoregions and biomes. A stronger decline in interaction diversity in the tropics and subtropics was also found in simulations including both obligate and opportunistic frugivores (Figure S4). In line with previous work, Neotropical species are sensitive to future climate change given their high degree of niche specialization and narrow climatic niches (Miranda et al., 2019). Many tropical species are also likely to occupy the warm edge of their climatic niche and will therefore be sensitive to future warming (Huey et al., 2009). Our findings are consistent with previous studies proposing that temperate bird species have a comparatively higher resilience because they experience a wider range of climate conditions across their larger geographic ranges (McCloy et al., 2022).

Our model projected declines in plant-frugivore interaction diversity across almost all major forest biomes. Declines in plant-frugivore interaction diversity were particularly pronounced in tropical and subtropical forest biomes, whereas the response of plant-frugivore networks in temperate biomes was more variable. The projected declines in plant-frugivore interaction diversity will likely reduce seed-dispersal functions in forest ecosystems, especially in the tropics, and can lead to the co-extinction of dependent plant species through the disruption of their life cycles, especially in many tropical plant species reliant on animals for seed dispersal (Jordano et al., 2011). Reduced interaction diversity can lead to lowered seed-dispersal distances in these assemblages (Donoso et al., 2020). The consequences of such effects are expected to be particularly accentuated in landscapes that have already suffered habitat degradation and whose functional connectivity is reliant on long-distance seed dispersal (Mueller et al., 2014). Forests will be particularly susceptible to such effects given the high proportion of plant species dependent on animal mediated seed dispersal in these ecosystems (Côrtes & Uriarte, 2013). The projected declines in plant-frugivore interaction diversity could therefore lead to a disruption of tropical and subtropical forest regeneration, driving cascading effects on other ecosystem functions and services such as carbon sequestration (Waring et al., 2020).

### Potential effects of interaction rewiring on interaction diversity

Integrating mechanisms of interaction rewiring between plants and birds compensated for projected future losses in interaction diversity and led to large increases in projected interaction diversity, especially in the tropics. Both mechanisms of dispersal and increased dietary flexibility would allow for new interactions to form between plants and frugivores (CaraDonna et al., 2017; Schleuning et al., 2020), either with new bird species entering an assemblage or with species already present. The simulations therefore suggest that these mechanisms can potentially buffer the projected negative impacts on plant-frugivore interaction diversity. However, these results have to be interpreted with caution as both rewiring scenarios are optimistic because they allowed for the formation of new interactions but did not incorporate mechanisms that could reduce future interaction diversity such as increased competitive interactions between frugivores (Lamb et al., 2017).

In simulations restricted to obligate frugivores, avian dispersal and increased dietary flexibility generated widespread positive effects on plant-frugivore interaction diversity across the Neotropics and in tropical and subtropical forest biomes. According to our findings, Neotropical assemblages would have a large potential for interaction rewiring because bird species from neighbouring assemblages could enter the networks and/or persisting species may be able to establish interactions to new plant partners. This pattern is likely driven by the large diversity of obligate frugivorous birds in the Neotropics. This diversity may facilitate the immigration of species into neighbouring ecoregions. Furthermore, the high trait diversity of Neotropical frugivores is likely to favour the formation of new interactions as especially large-gaped frugivores are able to interact with many different plant species (McFadden et al., 2022) and can potentially adapt their fruit diet under future conditions. Our simulations suggest that both mechanisms of interaction rewiring are additive because simulations incorporating both mechanisms together resulted in the largest increases in interaction diversity. For example, large-bodied frugivores, such as the red-ruffed fruitcrow (*Pyroderus scutatus* or the white-throated toucan (*Ramphastos tucanus)*, have a high dispersal capacity and therefore can reach a high number of other ecoregions, thus contributing to an increased interaction rewiring potential. Additionally, many such species have a large gape width and are therefore likely to have a high dietary flexibility in their prospective future assemblages (Bender et al., 2017).

According to our simulations of obligate frugivores, both interaction rewiring mechanisms had only weak positive effects on the Nearctic assemblages, likely due to the small species pool (Kissling et al., 2009) and the low diversity of obligate frugivores in these ecoregions (Dalsgaard et al., 2017; McFadden et al., 2022). The core of seed-dispersal networks composed by obligate frugivores is therefore likely to erode in such ecoregions under future climates. However, many frugivorous birds in temperate forest biomes have a generalist diet and are opportunistic frugivores (Dalsgaard et al., 2017; Schleuning et al., 2012; Belmaker et al., 2012). This explains why simulations including both obligate and opportunistic frugivores showed a buffering effect of interaction rewiring across both the Nearctic and Neotropics. The seed-dispersal functions of opportunistic frugivores may be crucial to buffer climatic effects on plant-frugivore interaction diversity especially in Nearctic ecoregions and temperate forest biomes. However, because many opportunistic species frequently switch between a fruit-based and an invertebrate diet (Carnicer et al., 2008), it is unknown to what extent these species may actually provide reliable seed-dispersal functions under future conditions. A key innovative element of our study was the incorporation of interaction rewiring through avian dispersal and increased dietary flexibility, both of which would have the potential to buffer the projected losses in interaction diversity under future conditions. As there is very little empirical evidence of how interaction rewiring may reorganize ecological assemblages (Kissling & Schleuning, 2015), it is difficult to judge the ecological realism of these projections. Because we employed trait-based approaches to model both dispersal and dietary flexibility, we are confident that our approach presents the best possible practice and goes beyond more simplistic approaches used in other studies (Schleuning et al., 2016; Richardson et al., 2000). Nevertheless, we acknowledge that our dispersal scenario only allowed for immigration of species but neglected that other species may go locally extinct due to competitive interactions with immigrating species (Thomas et al., 2006; Pigot et al., 2020). Similarly, a lower specificity of plant-bird interactions would likely lead to increased competition among bird species for fruit resources and could result in changes in interaction network structure. We therefore stress that a careful interpretation of these scenarios is needed as they were focussed on the potential beneficial and stabilizing effects of rewiring on interaction diversity, and it is unknown whether plant species will indeed experience widespread positive effects through interaction rewiring in future assemblages.

### Limitations of the simulations

We have generated macroscale projections of plant-frugivore interaction diversity by deploying a trait-based interaction model, informed by species-level functional trait and distribution data, coupled with climatic niche estimates. Key limitations of our study include spatiotemporal biases present in the underlying bird occurrence data used to generate the avian climate niche estimates (Johnston et al., 2019). Furthermore, while for birds there is comprehensive data on their geographic distributions and functional trait datasets, for plants these datasets are incomplete which spatially constrained this study and may have introduced some biases towards certain plant groups. Climatic niche estimates used in this study represent realised niche estimates which may underestimate fundamental tolerance. Additionally, the method does not control for potential species’ adaptations to new climatic conditions, and our simulations assumed that the avian climatic niches were constant under future conditions. It is therefore possible that adaptations of avian frugivores to novel climatic conditions could pose an additional buffer to maintaining plant-frugivore interaction diversity under future conditions.

## Conclusions

Using species traits to infer potential changes in plant-frugivore interactions, our simulations suggest a widespread vulnerability of plant-frugivore interactions to future climate change, especially in tropical and subtropical ecoregions and biomes. According to our analyses of interaction rewiring, the large diversity of traits and plant-frugivore interactions in Neotropical ecoregions is crucial to increase the resistance of tropical plant-frugivore networks to these future changes. Maintaining this diversity is therefore key to maintain seed-dispersal functions and tropical biodiversity in the future. Large-scale studies such as ours can identify how the composition of ecological assemblages may influence how plant-animal interactions respond to climate change, improving our knowledge on essential ecological processes and functions in the most vulnerable ecosystems and regions.

## Data accessibility statement

We sourced bird gape width measurements from McFadden et al., (2022); fruit length measurements from Dehling et al., (2016), Kissling et al., (2019) and Sinnott-Armstrong et al., (2018), bird geographical range data from BirdLife International (https://www.birdlife.org/), and plant geographical range data from BIEN (https://bien.nceas.ucsb.edu/bien/). All data and code for this study are deposited on Dryad (reviewer link): https://datadryad.org/stash/share/2j5QzuT14XjerbGUK-mKTBqhgUKYADPkeyPsRllkW3U.

## Conflict of interest statement

The authors declare no conflicts of interest.

## Notes

https://datadryad.org/stash/share/2j5QzuT14XjerbGUK-mKTBqhgUKYADPkeyPsRllkW3U

### Competing Interest Statement

The authors have declared no competing interest.

