## Supplementary Materials for "Projected impacts of climate change on plant-frugivore interactions across the Americas"

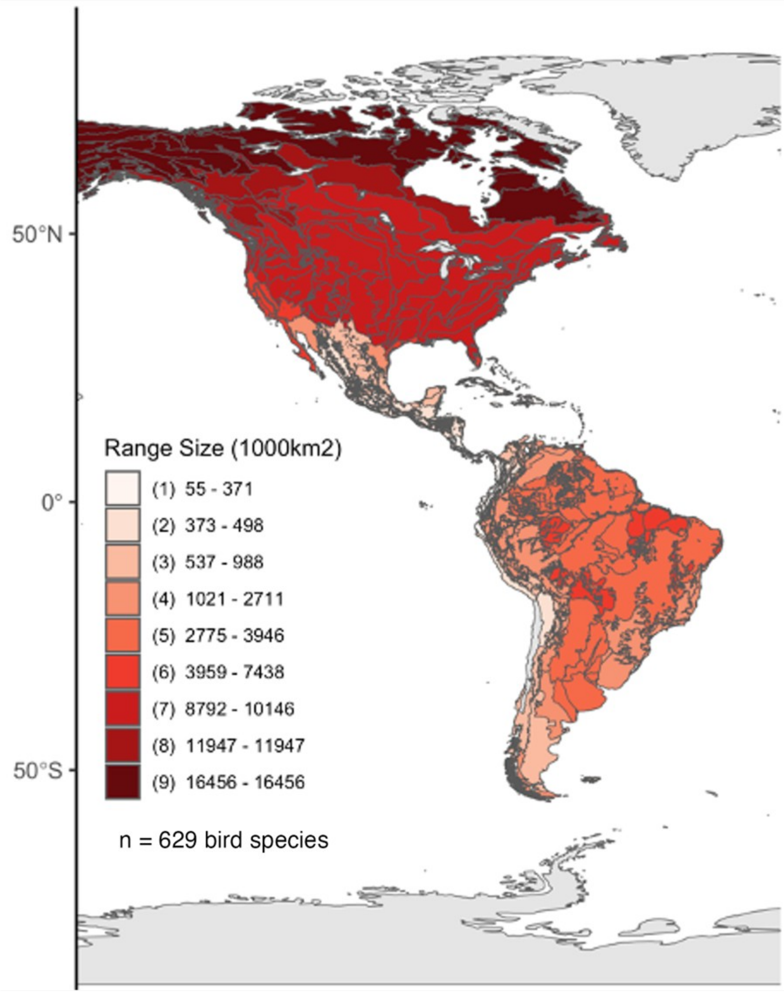


**Figure S1.** Map of median range size across bird species within an ecoregion. Darker red ecoregions are occupied by bird species that generally have larger ranges. Range size estimates of each bird species were extracted from the AVONET database and averaged across ecoregions.

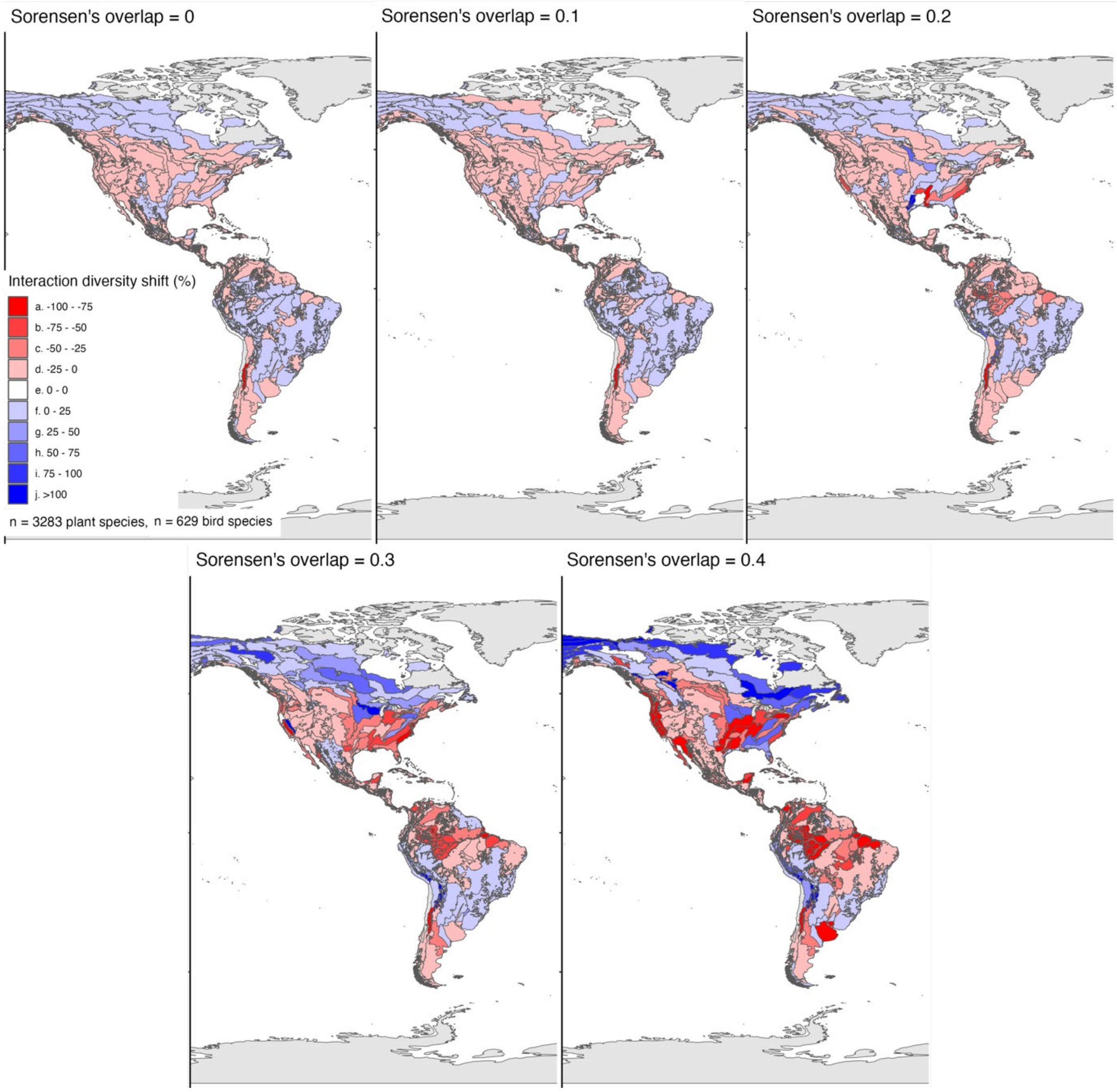


**Figure S2.** Maps of future climate impacts on plant-frugivore networks under different Sorensen's overlap threshold scenarios (Sorensen's overlap: 0, 0.1, 0.2, 0.3, 0.4). Effects sizes increase with an increase in the overlap threshold, but geographic patterns of changes in interaction diversity are qualitatively identical.

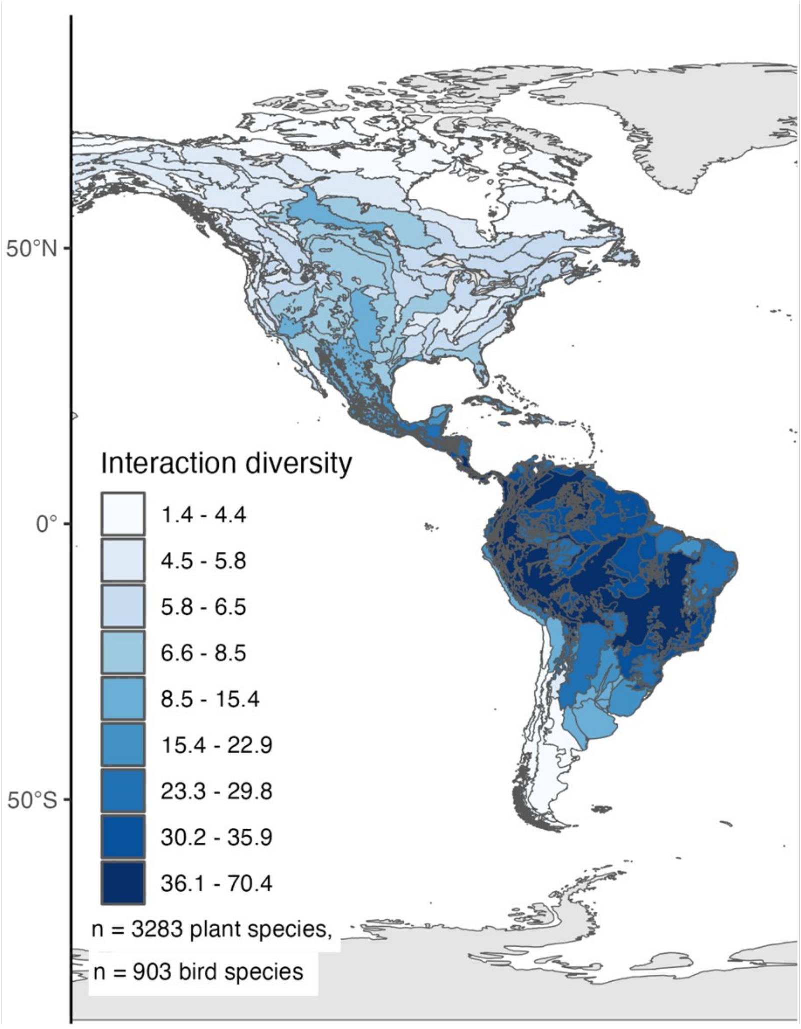


**Figure S3.** Map of the potential mean number of obligate and opportunistic avian frugivores per plant species within an ecoregion. Darker blue ecoregions are occupied by plant species that, on average, are able to interact with many more frugivorous birds compared to plant species that occupy lighter blue ecoregions. Plant interaction diversity shown here is not weighted by the climatic suitability of the respective ecoregions.

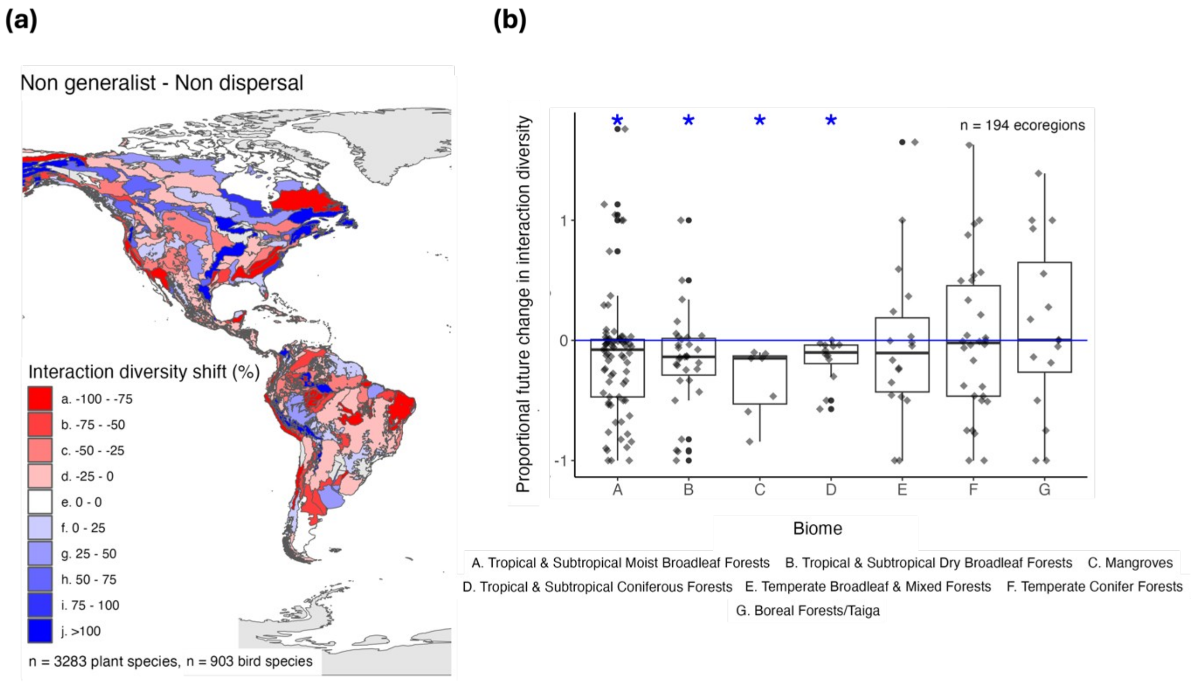


**Figure S4.** Climate change impacts on plant-frugivore networks including both opportunistic (>25% frugivory in their diet) and obligate frugivores (>50% frugivory in their diet). (a) Map of future climate impacts on plant-frugivore networks. (b) Boxplot of future climate impacts on plant-frugivore networks. Proportional change in interaction diversity was calculated by dividing future number of interaction partners by current number of interaction partners and substituting by 1. Asterisks above each boxplot indicate significant differences from 0 based on one-sample t-tests.

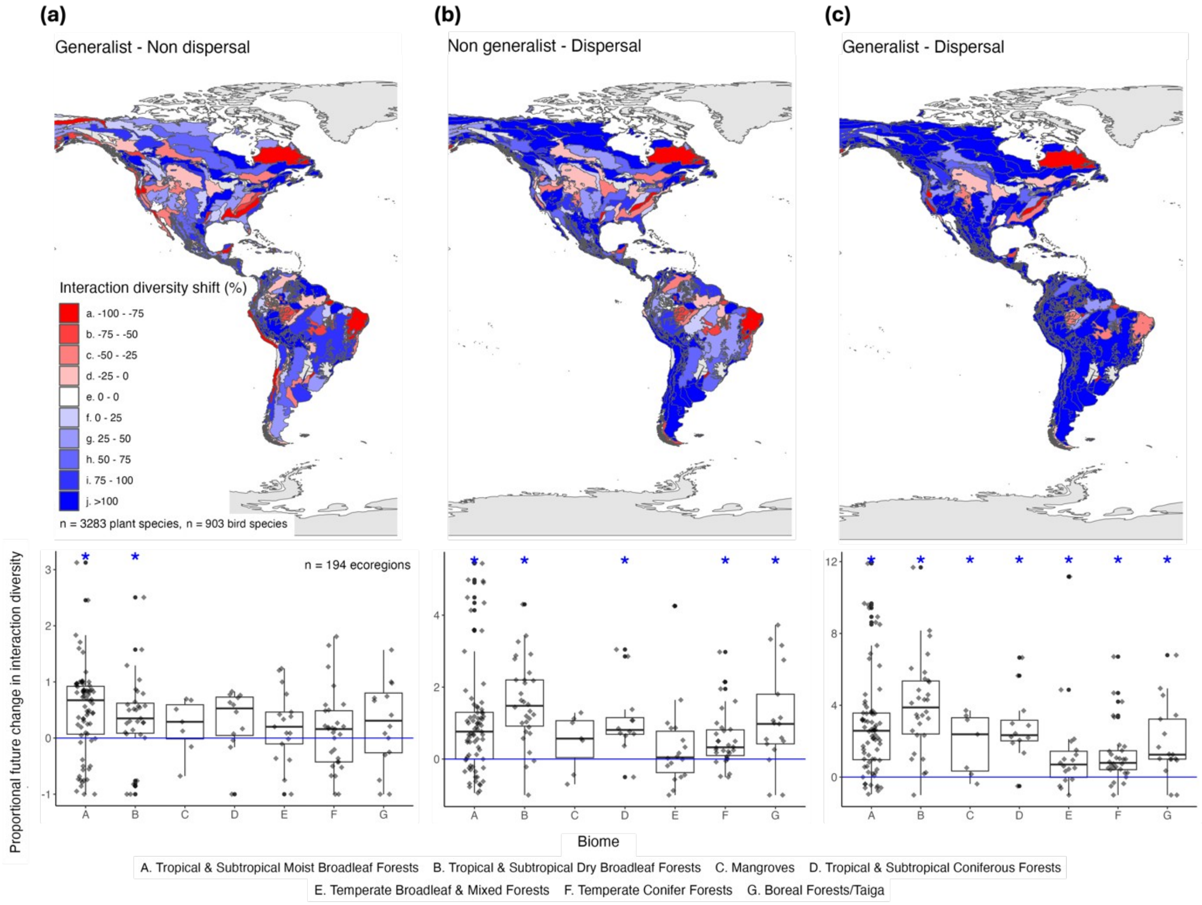


**Figure S5.** Climate change impacts on plant-frugivore networks including both opportunistic (>25% frugivory in their diet) and obligate frugivores (>50% frugivory in their diet) under different interaction rewiring scenarios. (a) Map and boxplot of future climate impacts on plant-frugivore networks allowing for avian dispersal (i.e., immigration of species from surrounding ecoregions). (b) Map and boxplot of future climate impacts on plant-frugivore networks allowing for dietary flexibility (i.e., trait matching parameter *s* set to 0.5). (c) Map and boxplot of future climate impacts on plant-frugivore networks allowing for both avian dispersal and dietary flexibility. Proportional chance in interaction diversity was calculated by dividing future number of interaction partners by current number of interaction partners and substituting by 1. Asterisks above each boxplot indicate significant differences from 0 based on one-sample t-tests. Note the different scaling on the y-axes for the different scenarios.
